# VenusGenome: Genome-Aware Masked Autoencoding for Microbial Genome Representation Learning

**DOI:** 10.64898/2026.10.24.761516

**Authors:** Liang Zhang, Yukun Zhang, Yang Tan, Fan Jiang, Bozitao Zhong, Xiaoran Cheng, Liang Hong, Mingchen Li, Yuanxi Yu

## Abstract

Genomic-context models have shown that contextualizing protein embeddings can improve predictions of microbial function. We observe that a gene’s protein embedding is partially predictable from the embeddings of neighboring genes in the genome, motivating high-ratio masking that requires reconstruction from sparser genomic context. Genomic modeling must also distinguish biological relationships from assembly conventions, since the ordering of separate contigs does not imply adjacency and strand labels depend on the reference orientation. We therefore introduce VenusGenome, a gene-resolution masked autoencoder that combines visible-only encoding with genome-aware attention incorporating contig boundaries, relative strand orientations, and gene positions. We pretrain this model on a curated corpus of over 1.7 million microbial genome assemblies from isolate and metagenomic sources, using frozen protein embeddings as gene-level inputs. Across genome-level phenotype prediction and gene-level tasks, the resulting frozen representations achieve the highest mean performance among the evaluated models on nine of eleven primary metrics. An exploratory gene-input perturbation analysis further links representation sensitivity to experimentally measured fitness defects. Ablations support the contribution of genome-aware attention and show the benefits of masked autoencoding over BERT-style masked modeling, with further gains from higher masking ratios.

## 1 Introduction

Microbial function depends on both the proteins a genome encodes and the contexts in which their genes occur. Gene order, orientation, and co-occurrence organize proteins into operons, biosynthetic regions, and other functional modules (Taboada et al., 2018; Hannigan et al., 2019; Carroll et al., 2021). Learning representations that retain this organization could support predictions from individual-gene function to organismal phenotype.

Protein language models provide transferable molecular representations, but independent protein encoding omits genomic organization (Rives et al., 2021; Lin et al., 2023). Nucleotide models retain sequence context at the cost of processing millions of bases per microbial assembly. Gene-context models offer an intermediate resolution: gLM contextualizes protein embeddings in short metagenomic segments, and BacFormer extends ordered protein modeling to bacterial genomes (Hwang et al., 2024; Wiatrak et al., 2025a). Their pretraining targets differ: gLM predicts continuous protein features, while BacFormer predicts discrete protein-family labels. These models show that contextualizing protein embeddings can yield useful gene representations, while both use 15% masking and retain masked positions in BERT-style encoders during pretraining.

However, sparse masking can leave substantial local context available for reconstruction. In a preliminary analysis of our pretraining embeddings, a linear predictor using the two preceding and two following genes reduced reconstruction error by 15.4% relative to a constant mean predictor, compared with 8.9% using four random genes from the same contig. This result indicates that local neighborhoods provide readily accessible reconstruction cues beyond contig-level composition. In vision, the availability of local reconstruction cues motivates high-ratio masking in masked autoen-coders (He et al., 2022). Although local predictability in protein embeddings differs from spatial redundancy in images, our observation motivates higher masking ratios, which reduce the availability of immediate neighbors and require reconstruction from sparser genomic context.

Genomic organization introduces a complementary modeling challenge: distinguishing biological relationships from arbitrary assembly conventions. Existing models already incorporate structural information, including gene orientation in gLM and contig embeddings and rotary positional encodings in BacFormer (Hwang et al., 2024; Wiatrak et al., 2025a). However, the ordering of separate contigs does not establish genomic adjacency, and strand labels depend on the chosen reference orientation. The same- or opposite-strand relationship between genes within a contig, in contrast, is preserved when that orientation is reversed. These considerations motivate explicitly distinguishing within-contig interactions from cross-contig communication and encoding pairwise strand relationships rather than relying solely on individual strand labels.

To address these limitations, we introduce VenusGenome, a microbial genome model that learns contextualized gene and whole-genome representations from frozen protein embeddings. We represent each genome as an ordered sequence of gene-level protein features and train an asymmetric masked autoencoder to reconstruct missing features from genomic context. Motivated by the observed predictability of protein embeddings from neighboring genes, we use 75% masking to require reconstruction from sparser genomic context. The encoder processes only visible genes, while a lightweight decoder reconstructs the masked embeddings, separating contextual encoding from reconstruction and reducing encoder computation during pretraining. To account for genomic organization, we combine contig-restricted gene attention with relative strand biases and within-contig positional information. Global tokens mediate communication across contigs without treating their concatenation as a continuous genomic neighborhood. We pretrain VenusGenome on a curated corpus of over 1.7 million microbial genome assemblies and evaluate the frozen encoder on genome-level phenotype prediction and gene-level functional tasks. We further examine whether sensitivity to gene-input perturbation can identify genes with experimentally measured average fitness defects. Our main contributions are as follows:

- We develop a genome-aware masked autoencoding framework that combines visible-only contextual encoding with structured genomic attention, and evaluate its main design choices through controlled ablations. At 512M and matched 15% masking, MAE reaches a zero-shot protein–protein interaction (PPI) genome-macro Spearman correlation of 0.282 versus 0.192 for BERT-style reconstruction; removing contig-aware attention from the 75%-masking model reduces it from 0.293 to 0.203. At this scale, increasing masking from 15% to 75% also improves correlation from 0.282 to 0.293 and reduces measured forward–backward time by 65.2% for 4,096-gene inputs.
- We curate over 1.7 million isolate and metagenomic microbial genome assemblies into a unified gene-resolution pretraining corpus. Quality filtering, taxonomic standardization, and cross-source annotation integration preserve gene order, contig membership, and strand information, enabling genome-aware pretraining at scale.
- We evaluate one frozen encoder on six whole-genome phenotype tasks and five gene-level metrics spanning biosynthetic gene clusters (BGCs), essential genes, operons, and PPI. VenusGenome leads evaluated models on nine of eleven primary metrics, including zero-shot PPI ranking (0.300 versus 0.201 for BacFormer). An exploratory zero-shot analysis further links representation sensitivity to experimentally measured gene-disruption fitness, suggesting a potential route toward in silico functional screening in microbes.

## 2 Related Work

### Biological sequence and genome-context models

Protein language models learn transferable molecular representations (Alley et al., 2019; Elnaggar et al., 2022; Rives et al., 2021; Meier et al., 2021; Lin et al., 2023), with alignment-aware, generative, and structure-aware extensions (Rao et al., 2021; Madani et al., 2023; Su et al., 2024b). Nucleotide models learn from coding and non-coding sequences through masked, autoregressive, and long-context architectures (Ji et al., 2021; Zhou et al., 2024; Dalla-Torre et al., 2025; Zvyagin et al., 2023; Nguyen et al., 2023; Schiff et al., 2024; Nguyen et al., 2024; Brixi et al., 2026; Fang et al., 2026). At gene resolution, gLM contextualizes protein embeddings in metagenomic segments and predicts continuous protein features (Hwang et al., 2024), whereas BacFormer models ordered protein embeddings across bacterial genomes with protein-family prediction targets (Wiatrak et al., 2025a). FGeneBERT combines protein-based gene representations with masked gene modeling and contrastive learning for metagenomics (Duan et al., 2025). Other approaches represent genomes through mixed coding and intergenic sequences, as in gLM2 (Cornman et al., 2025), or protein sets, as in Protein Set Transformer (Martin et al., 2025).

### Masked representation learning and structured attention

BERT and BART learn representations by predicting masked or corrupted inputs (Devlin et al., 2019; Lewis et al., 2020). MAE instead encodes only visible inputs and uses a lightweight decoder for reconstruction, allowing high masking ratios to reduce encoder computation (He et al., 2022). MAE-LM extends masked autoen-coding to language modeling and shows that excluding mask tokens from the encoder can improve the utilization of representation dimensions by real tokens (Meng et al., 2024). Complementing this architectural choice, MaskFeat predicts fixed features and I-JEPA predicts learned target representations, providing alternatives to raw-input reconstruction (Wei et al., 2022; Assran et al., 2023). Structured attention controls which inputs interact and how their relationships affect attention: Big-Bird combines local, random, and global connections (Zaheer et al., 2020), while T5 introduces learned per-head relative-position biases (Raffel et al., 2020).

## 3 Method

VenusGenome first maps frozen protein representations and assembly metadata to an ordered gene sequence, then learns contextualized states by reconstructing genes removed from a visible-only encoder. The same encoder is used without masking for downstream gene- and genome-level representation extraction. Figure 1a summarizes the overall workflow, and Figure 1b previews the genome-aware attention mechanisms described below.

**Figure 1:**
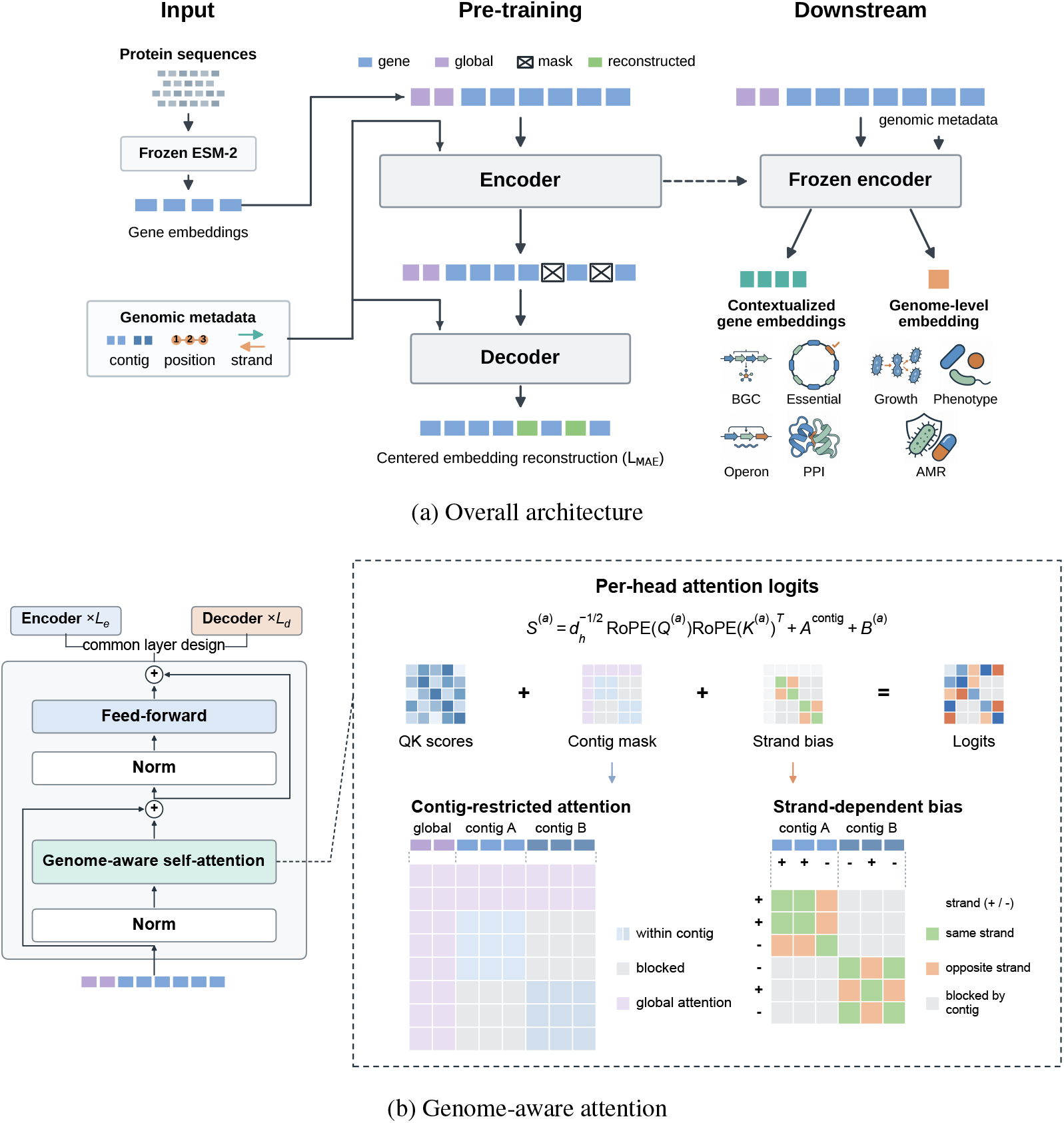
VenusGenome architecture and attention design. (a) Frozen protein embeddings and genomic metadata form gene inputs; a visible-only encoder and lightweight decoder perform masked reconstruction, and the frozen encoder supports downstream tasks. (b) Encoder and decoder use the same layer design with separate parameters. Contig masks and strand biases modify RoPE query– key logits; the expanded matrices show global connectivity and within-contig gene relationships.

### 3.1 Inputs and Genome-aware Attention

Let a microbial genome be an ordered sequence of *n* protein-coding genes, *G* = (*g*_1_, …, *g*_*n*_). Each gene *g*_*i*_ has a 480-dimensional ESM2-35M embedding ***x***_*i*_ ∈ ℝ^*d*^ from the frozen protein language model (Lin et al., 2023), obtained by averaging residue states from its 12th (final) Transformer layer while excluding special and padding tokens, together with metadata ***m***_*i*_ = (*c*_*i*_, *p*_*i*_, *s*_*i*_) denoting its contig, within-contig ordinal position, and strand. We project the raw protein embedding as ***e***_*i*_ = *W*_*x*_***x***_*i*_ + ***b***_*x*_. A per-dimension corpus mean ***µ***, estimated over the pretraining protein embeddings, is used only to center the reconstruction targets; encoder inputs are not divided by a per-dimension standard deviation.

After selecting the visible genes, we prepend *G* learned global tokens to the encoder sequence. They attend bidirectionally to every visible gene and to one another, aggregating genome-wide information and relaying information between contigs that otherwise cannot communicate directly. The resulting global states are projected into the decoder together with the visible-gene states, where they retain global connectivity to both visible and masked positions.

Metadata modifies self-attention directly. We apply RoPE (Su et al., 2024a) using positions *π*_*i*_ = *p*_*i*_ + *δ* for gene tokens and *π*_*i*_ = 0 for global tokens. During pretraining, one offset *δ* is sampled uniformly from {0, …, 2047} per input window and shared across its genes and both encoder and decoder; at inference, *δ* = 0. For attention head *a*, let 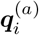 and 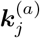 denote query and key vectors of width *d*_*h*_; the pre-softmax logit is

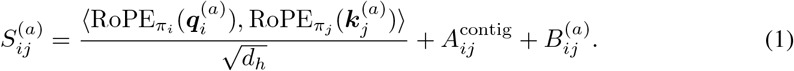

Here 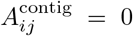 when *i* and *j* belong to the same contig or either token is global, and −∞ otherwise. For gene pairs within the same contig, 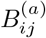 equals the learned head-specific scalar 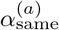 or 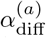 according to whether their strands agree; it is zero for all other pairs. The encoder and decoder learn separate strand-bias parameters, each shared across the layers of its respective module. Figure 1b illustrates how these components enter each Transformer layer and determine gene connectivity.

### 3.2 Masked Feature Reconstruction

VenusGenome follows an asymmetric Transformer masked autoencoder (Vaswani et al., 2017; He et al., 2022). Pretraining uses at most 4,096 genes per example, with a contiguous gene window randomly cropped from longer assemblies. We uniformly sample 75% of valid genes to form the masked set ℳ, leaving the visible set *V* = {1, …, *n*}\ ℳ. The encoder processes only the visible gene embeddings and global tokens, preserving the genes’ original within-contig positions. The lightweight decoder projects the encoder states, inserts a learned mask token at each masked gene position, and restores the original gene order. It retains the global tokens and uses the same positional, contig, and strand mechanisms as the encoder, with separate learned parameters, to produce gene states 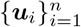 .

We center the protein reconstruction targets to reduce the contribution of shared embedding directions to cosine similarity, motivated by anisotropy observed in language-model representations (Ethayarajh, 2019). In a diagnostic using ESM2 embeddings from 1,000 genomes, subtracting the fixed pretraining mean reduces mean cosine similarity between random cross-genome protein pairs from 0.830 to 0.012. With targets ***y***_*i*_ = ***x***_*i*_ − ***µ*** and predictions ***ŷ***_*i*_ = *r*_*ϕ*_(***u***_*i*_), we minimize reconstruction loss only at masked gene positions:

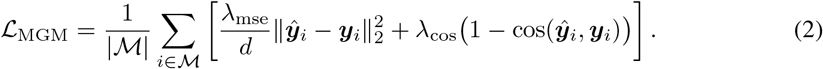

We set *λ*_mse_ = 0.3 and *λ*_cos_ = 0.7, combining Euclidean reconstruction with angular alignment. The canonical model pairs a 44-layer encoder with an 8-layer decoder; Appendix A.4 provides full architecture and optimization settings.

### 3.3 Representation Extraction

For downstream extraction, we set the mask ratio to zero, discard the decoder, and encode every valid gene. Gene-level tasks use the resulting contextualized state ***h***_*i*_ directly; genome-level tasks use either the mean of all gene states or the mean of the final encoder global states,

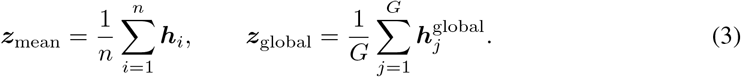

For assemblies longer than 4,096 genes, we use overlapping 4,096-gene windows with 512-gene overlap, average repeated gene states across windows, and average the window-level global readouts. Windows are encoded independently, so stitching covers every gene but does not introduce direct attention across windows. Main genome results use ***z***_mean_ from the stitched gene states. Appendix B.1 compares both readouts on the same species-disjoint benchmarks.

## 4 Experiments

We evaluate VenusGenome across genome- and gene-level benchmarks, covering six whole-genome phenotype tasks, BGC detection and category prediction, essential-gene prediction, and zero-shot operon and PPI ranking. We also assess zero-shot gene fitness prediction using experimental measurements from Fitness Browser.

### 4.1 Experimental Setup

#### Pretraining setup

We assemble a pretraining corpus of over 1.7 million microbial genomes from NCBI, proGenomes v4, MGnify, and SPIRE (Sayers et al., 2022; Fullam et al., 2026; Gurbich et al., 2023; Schmidt et al., 2024). For isolate genomes, we limit overrepresented species and apply completeness and contamination filters, retaining 75,390 high-quality assemblies. Unless otherwise specified, VenusGenome denotes the 1B encoder pretrained for 40k steps with 75% gene masking and contexts of up to 4,096 genes; 85M and 512M variants are used for scaling analysis. Appendices A.1 and A.4 detail corpus construction and pretraining settings.

#### Downstream evaluation

We compare ESM2 protein embeddings (Lin et al., 2023), gLM and BacFormer gene-context embeddings (Hwang et al., 2024; Wiatrak et al., 2025a), and Nucleotide Transformer v2 (NT-v2) and Evo2 embeddings (Dalla-Torre et al., 2025; Brixi et al., 2026). For genome-level tasks, both nucleotide baselines aggregate features extracted from DNA windows across each assembly, including coding and intergenic regions; VenusGenome uses mean pooling of contextualized gene states. All representation encoders remain frozen. The six supervised genome-level tasks use species-disjoint splits, while BGC and essential-gene prediction use genome-disjoint splits. Within each task, we use the same one-hidden-layer MLP probe, optimization settings, fixed data split, and validation-based checkpoint selection across representations, adjusting input dimensions to the embedding size. Supervised results are averaged over five probe-training seeds; operon, PPI, and gene fitness evaluations are zero-shot. Appendices A.2–A.4 provide evaluation and extraction details.

### 4.2 Genome-Level Evaluation

We evaluate genome-level representations on six supervised tasks covering microbial growth traits (optimal growth temperature and population doubling time), antimicrobial response (MIC regression and multilabel AMR classification), and binary and categorical microbial phenotypes. To assess transfer to unseen species, we use species-disjoint training, validation, and test sets defined by NCBI species taxon identifiers. Table 1 summarizes the results; task-specific metrics and evaluation protocols are detailed in Appendix A.2.

**Table 1:** Genome-level supervised results under strict species-disjoint splits.

| Model | *(a) Regression* |  |  | *(b) Classification* |  |  |
| --- | --- | --- | --- | --- | --- | --- |
|  | Temperature<br>Pearson | Doubling<br>Pearson | MIC<br>Macro Pearson | AMR<br>Macro AUROC | Binary phen.<br>Macro AUROC | Mixed phen.<br>Macro F1 |
| ESM2-35M | 0.845 ± 0.005 | 0.212 ± 0.016 | 0.110 ± 0.041 | 0.520 ± 0.018 | 0.771 ± 0.004 | 0.420 ± 0.025 |
| ESM2-650M | **0.877** ± 0.002 | 0.217 ± 0.010 | 0.197 ± 0.038 | 0.574 ± 0.010 | 0.783 ± 0.003 | 0.389 ± 0.025 |
| gLM | 0.848 ± 0.004 | 0.072 ± 0.146 | 0.175 ± 0.019 | 0.560 ± 0.022 | 0.780 ± 0.006 | 0.410 ± 0.022 |
| BacFormer | 0.824 ± 0.001 | 0.185 ± 0.011 | 0.254 ± 0.015 | 0.614 ± 0.018 | 0.788 ± 0.001 | 0.421 ± 0.023 |
| NT-v2 | 0.755 ± 0.005 | 0.164 ± 0.047 | 0.153 ± 0.024 | 0.633 ± 0.026 | 0.756 ± 0.008 | 0.367 ± 0.011 |
| Evo2 | 0.476 ± 0.009 | 0.254 ± 0.005 | 0.152 ± 0.021 | 0.555 ± 0.063 | 0.687 ± 0.006 | 0.266 ± 0.014 |
| VenusGenome | 0.862 ± 0.001 | **0.336** ± 0.011 | **0.270** ± 0.016 | **0.682** ± 0.034 | **0.794** ± 0.004 | **0.457** ± 0.027 |

VenusGenome achieves the highest mean performance on five of six tasks under species-disjoint evaluation. The strongest competing baseline varies across traits: ESM2-650M leads on temperature, Evo2 on doubling time, NT-v2 on AMR, and BacFormer on MIC and both phenotype tasks. The gains therefore extend across traits favored by different representation models, with doubling-time Pearson correlation increasing from 0.254 for Evo2 to 0.336. VenusGenome also consistently outperforms mean pooling of ESM2-35M embeddings, which serve as its frozen inputs, showing that genome-level pretraining improves transfer beyond simply aggregating individual-protein representations.

### 4.3 Gene-Level Evaluation

We evaluate gene representations on supervised BGC and essential-gene prediction, together with zero-shot operon and PPI ranking. Zero-shot operon and PPI ranking further tests whether the learned representation geometry reflects functional relationships between genes, without a trained prediction head. Supervised evaluations use genome-disjoint splits; BGC detection includes all labeled genes, whereas category prediction is restricted to BGC-positive genes. Both zero-shot tasks use gene-embedding cosine similarity, with PPI performance measured by its Spearman correlation with STRINGDB confidence scores (Szklarczyk et al., 2023) within each genome and averaged equally across genomes. Table 2 summarizes the results; detailed evaluation protocols are provided in Appendix A.2.

**Table 2:**
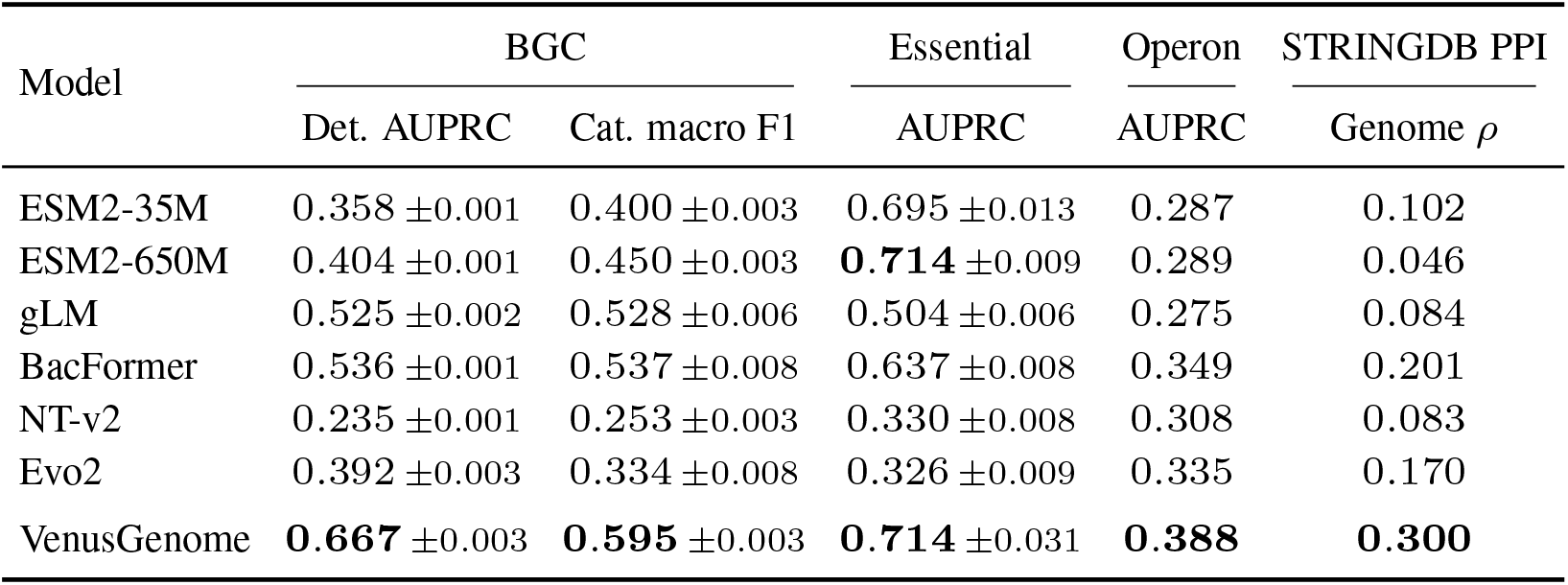
Gene-level results.

Table 2 shows that VenusGenome leads four of five metrics and performs comparably to ESM2-650M on essential-gene prediction. It improves on ESM2-35M, which supplies its frozen input embeddings, across all five metrics, demonstrating the value of additional genome-level pretraining. Notably, increasing ESM2 from 35M to 650M improves supervised performance but does not consistently improve zero-shot ranking. This distinction highlights that features useful to a trained predictor need not encode functional relationships directly through cosine similarity. VenusGenome improves both supervised prediction and zero-shot ranking, with PPI genome-macro Spearman correlation reaching 0.300, compared with 0.201 for BacFormer, the strongest competing baseline. Section 5.2 further examines how extraction context length affects BGC prediction and zero-shot PPI ranking.

### 4.4 Zero-Shot Gene Fitness Evaluation

We ask whether sensitivity to gene-input perturbation can identify genes associated with negative average fitness effects across assayed experiments. Fitness Browser 2024 provides gene fitness measurements from 7,552 pooled transposon-insertion experiments across 48 organisms (Price & Deutschbauer, 2024). Fitness reflects normalized log_2_ changes in mutant abundance, with negative values indicating a competitive disadvantage under the tested conditions. We hypothesize that genes with greater influence on neighboring representations, or higher sequence likelihood under Evo2, are more likely to be functionally important, such that their disruption leads to larger fitness defects.

Without fitting a downstream predictor, we replace each target gene’s protein embedding with its organism’s mean embedding and measure the change in the mean contextual representation of nearby genes. We label genes with mean experimental fitness ≤ − 1 as positive and all remaining evaluated genes as negative, and compute AUROC separately within each organism. VenusGenome achieves a median AUROC of 0.673, compared with 0.631 for an exploratory Evo2 baseline using standalone CDS likelihood, with higher AUROC in 39 of 48 organisms (Figure 2). These results suggest that local representation sensitivity carries information about experimentally measured average fitness defects. Appendix A.3 details the experimental targets, sensitivity scores, and evaluation protocol.

**Figure 2:**
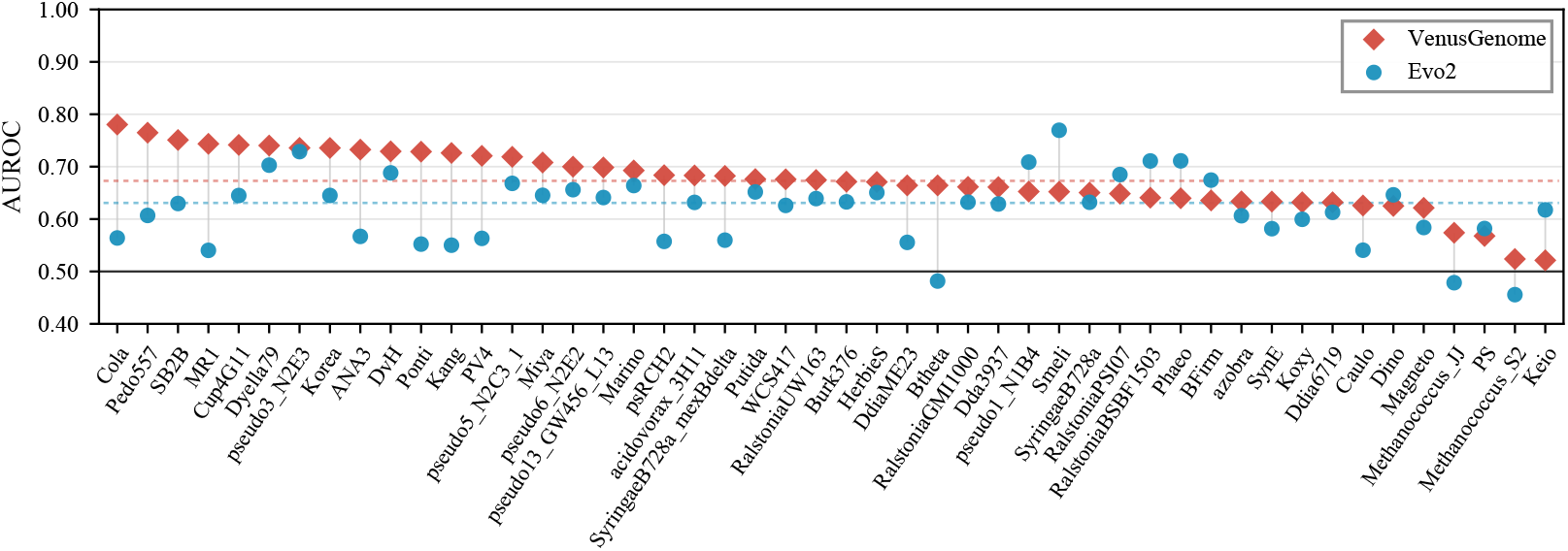
Zero-shot gene fitness evaluation on Fitness Browser 2024. Positive genes have mean experimental fitness ≤ − 1. Points show within-organism AUROC for all 48 organisms, ordered by VenusGenome; gray segments pair models and colored dashed lines mark medians. The black line marks random ranking (AUROC = 0.50).

## 5 Analysis

### 5.1 Pretraining Design

We analyze encoder scaling and conduct ablation studies on masking ratio, genome-aware attention, and pretraining formulation. We use zero-shot PPI ranking to test whether functional relationships between genes are reflected in the learned representation geometry. By comparing embedding cosine similarity with STRINGDB confidence scores, this evaluation tracks changes across checkpoints without fitting a downstream predictor. Figure 3 summarizes these comparisons.

**Figure 3:**
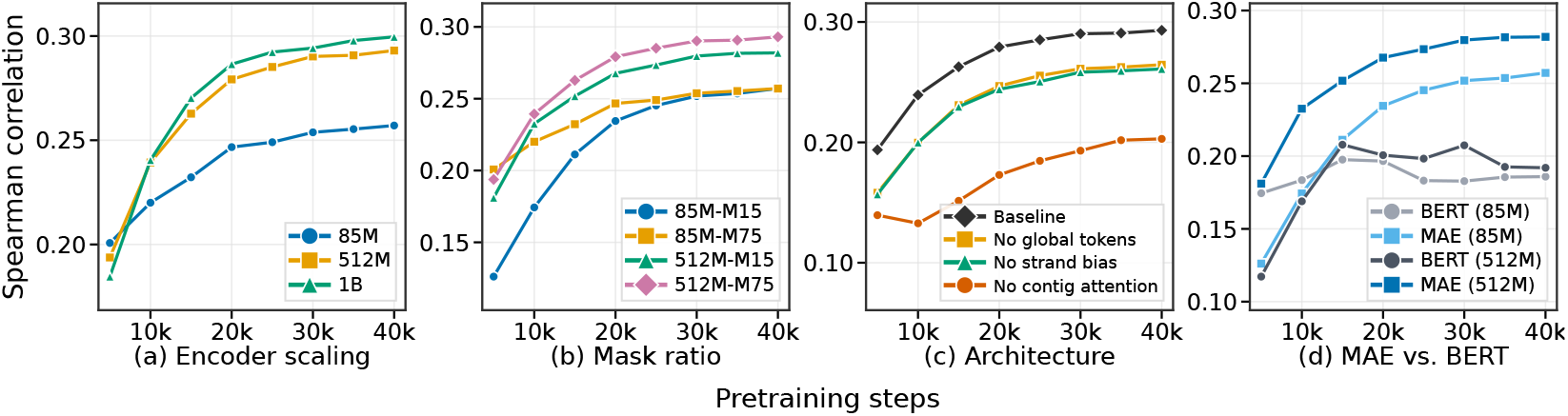
PPI scaling and design analysis. All panels report zero-shot genome-macro Spearman correlation on STRINGDB. (a) Encoder scaling at 75% masking. (b) Mask-ratio comparison. (c) Architectural ablations for the 512M model. (d) MAE versus BERT-style full-sequence reconstruction at 15% masking. Curves show single pretraining runs.

Increasing encoder size improves zero-shot PPI ranking under 75% masking, although the gain from 512M to 1B is smaller than that from 85M to 512M (Figure 3a). Higher masking does not uniformly improve ranking: moving from 15% to 75% improves the 512M model but leaves the 85M result nearly unchanged (Figure 3b). For the benchmarked 4,096-gene configuration, 75% masking reduces measured forward–backward time by 65.2% and peak memory by 51.3% relative to 15% masking (Appendix B.3). These results complement the local-predictability analysis in the Introduction: although neighboring genes provide readily accessible reconstruction cues, reducing their availability during pretraining preserves or improves PPI ranking in the evaluated configurations while substantially reducing computation.

Architectural ablations show that removing contig-aware attention causes the largest PPI degradation among the tested components (Figure 3c). At 512M, removing it reduces correlation from 0.293 to 0.203; removing global tokens or strand bias produces smaller declines. The pretraining comparison further shows that MAE outperforms BERT-style full-sequence reconstruction at both evaluated scales under 15% masking (Figure 3d). Both formulations reconstruct the same continuous targets with the same loss, but MAE separates visible-gene encoding from masked-gene reconstruction through a lightweight decoder. Appendix B.2 provides complementary pretraining-loss diagnostics.

We next examine whether MAE and BERT-style models differ in how broadly variation is distributed across their gene-representation spaces, following diagnostics in MAE-LM (Meng et al., 2024). We compare 85M and 512M checkpoints pretrained for 40k steps with 15% masking. For each of three genome-sampling seeds, we sample 256 isolate-source pretraining genomes and extract final normalized gene representations from the same un-masked windows of up to 512 genes. We select up to 64 identical gene positions per window for both models, excluding global tokens. For each model, we subtract the mean of all sampled gene representations and use PCA to count the components needed to explain 90% of the variance. MAE requires more components than BERT at both scales (Figure 4), showing that variation among its gene representations is less concentrated in a small number of directions.

**Figure 4:**
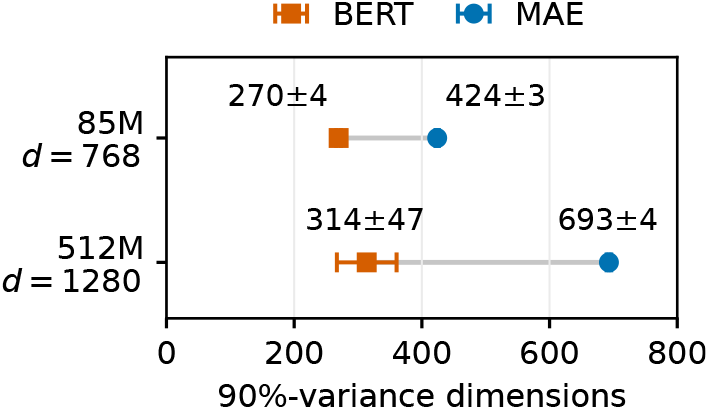
Centered gene-representation dimensionality (mean *±* SD over three genome-sampling seeds).

### 5.2 Context at Inference

We next ask how much genomic context is needed when extracting representations from a fixed pretrained model. Using the same 1B mask-75% checkpoint, we vary the extraction window from 9 to 4,096 genes and evaluate BGC prediction and zero-shot PPI ranking. The *single gene + global* control retains the pretrained encoder and all 64 global tokens but encodes each gene independently, removing access to other gene inputs. For a controlled comparison across context lengths, all BGC settings use the same fixed genome-disjoint split and five probe seeds as Table 2. PPI uses the same test pairs and genome-macro Spearman correlation as the main evaluation.

Figure 5 shows that both tasks benefit from additional gene context. A 33-gene window recovers approximately 90% of the improvement over the single-gene control observed at the maximum window size for both metrics. The benefit of further expansion differs between tasks: BGC AUPRC continues to increase, from 0.641 at 33 genes to 0.667 at 4,096 genes, whereas PPI correlation shows diminishing gains and is nearly unchanged beyond 513 genes. Thus, local neighborhoods account for much of the observed context benefit, while larger windows provide additional, more modest gains for BGC prediction.

**Figure 5:**
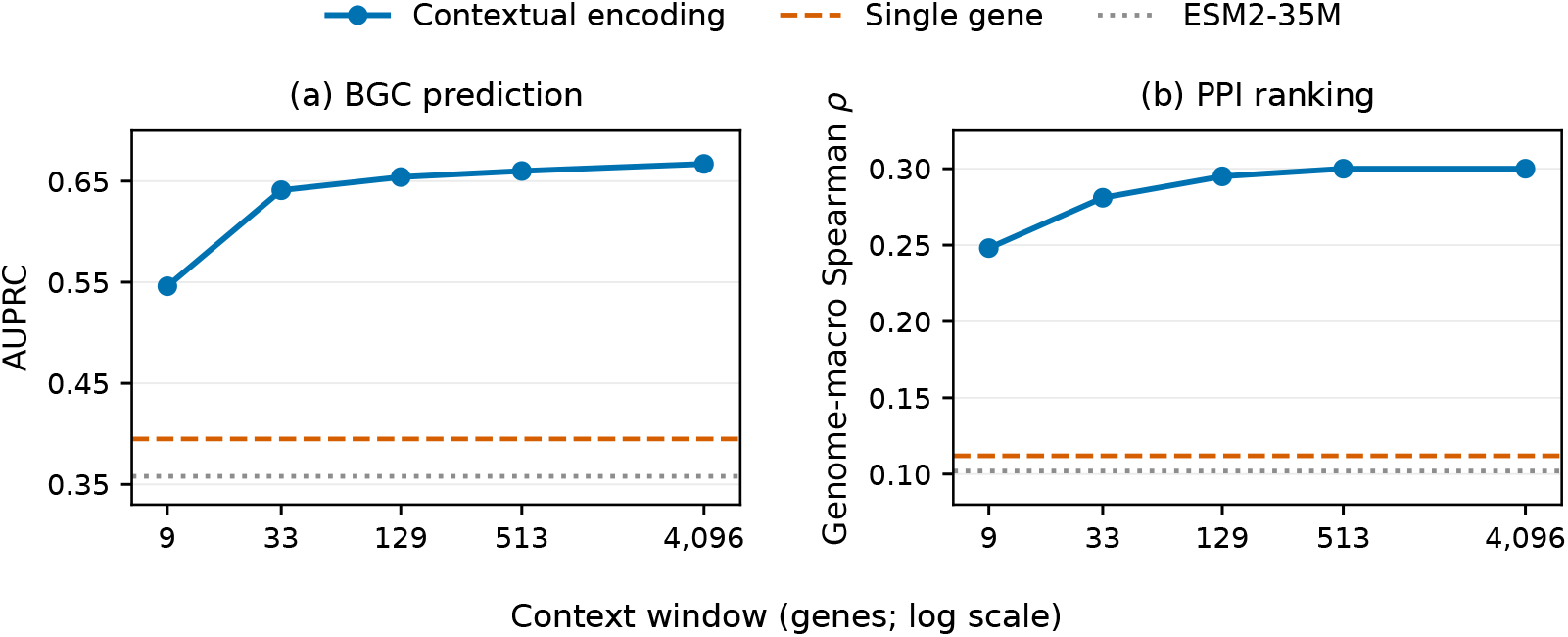
Effect of extraction context length with a fixed pretrained encoder. BGC reports mean AUPRC over five probe seeds on a fixed genome-disjoint split; PPI reports genome-macro Spearman correlation. Horizontal lines show the single-gene control and ESM2-35M baseline. Context length is plotted on a logarithmic axis.

## 6 Conclusion

We introduce VenusGenome, a microbial genome model that combines high-ratio masked autoen-coding with genome-aware attention to learn contextualized gene and whole-genome representations from protein embeddings. Pretrained on a curated corpus of over 1.7 million microbial genome assemblies, VenusGenome achieves the highest mean performance among the evaluated models on nine of eleven primary metrics using frozen representations. These results support its use as a shared representation model for microbial functional prediction at both gene and genome levels.

This study has several limitations that motivate further work. First, the use of frozen protein embeddings constrains contextual learning to the information retained by the protein encoder. Stronger protein representations, joint training, or complementary nucleotide information could expand the information available to the model. Second, our exploratory gene-fitness analysis evaluates average effects across assayed experiments and does not resolve condition-specific responses. Integrating genomic representations with environmental context and condition-specific perturbation measurements could extend this framework toward predicting how gene function and perturbation effects vary across conditions.

## Code Availability

The GitHub repository and Hugging Face model links will be made public when the paper is published online.

## A Experimental Details

### A.1 Datasets

The pretraining corpus combines isolate genomes from NCBI and proGenomes v4 (Sayers et al., 2022; Fullam et al., 2026) with metagenome-assembled genomes (MAGs) from MGnify and SPIRE (Richardson et al., 2023; Gurbich et al., 2023; Schmidt et al., 2024). Isolate taxonomy was standardized against LPSN (Parte et al., 2020), and the collection was capped at five genomes per species. Only the 75,390 high-quality isolate genomes passing CheckM (Parks et al., 2015) thresholds of ≥ 90% completeness and ≤ 5% contamination were included in the pretraining corpus. Genes were predicted with Prodigal (Hyatt et al., 2010) for isolates and taken from source annotations for MAGs; gene order, contig, and strand were retained. The MAG collection contains more than 1.67 million genomes, with more than 510k from MGnify and 1.16 million from SPIRE. Figure 6 compares their phylum-level composition after species-level dereplication.

**Figure 6:**
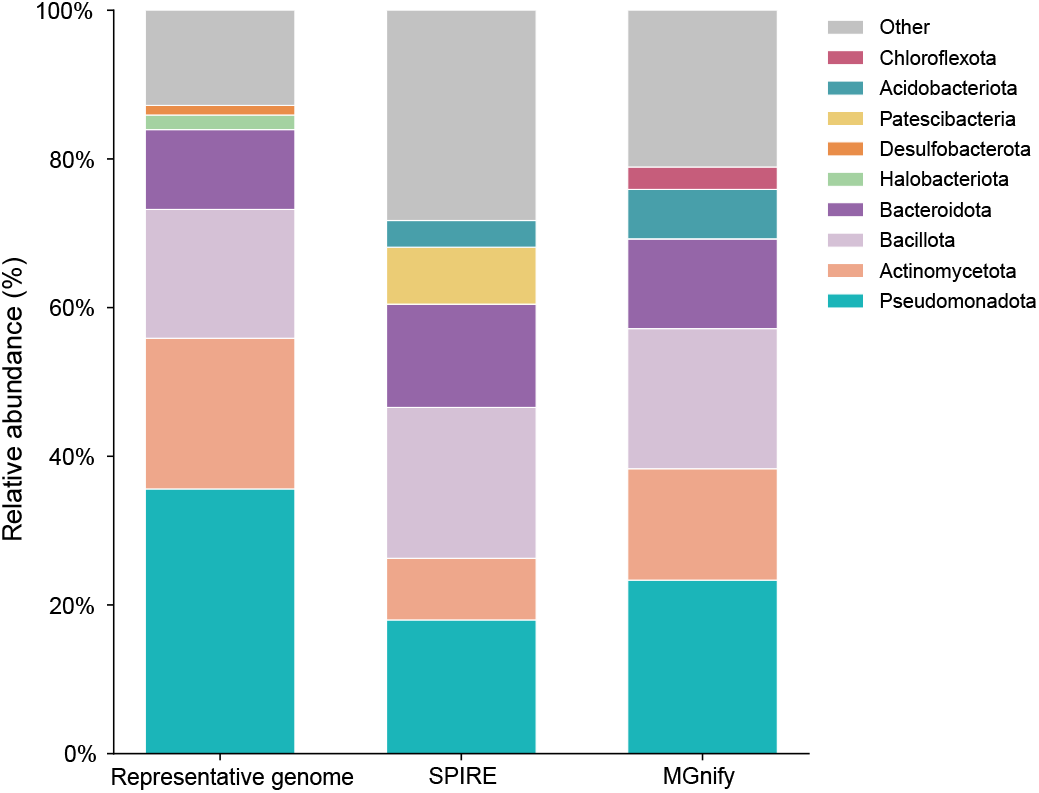
Phylum-level composition of the representative high-quality isolate collection, SPIRE, and MGnify at the species-cluster level. Each stacked bar reports the relative abundance of non-redundant species clusters within one source. The legend is the union of the six most abundant phyla selected separately in each source; phyla outside a source-specific top six are grouped as *Other* for that source. Each species cluster is counted once within a source, and phylum labels are harmonized across the source taxonomies.

Genome-level benchmarks cover six tasks: optimal growth temperature, doubling time, binary antimicrobial resistance, minimum inhibitory concentration (MIC), binary phenotypes, and mixed phenotypes, using data derived from BacDive, BacBench, and BacFormer (Reimer et al., 2022; Wiatrak et al., 2025b;a). All six main genome-level results use species-disjoint splits based on NCBI species taxon identifiers, with no species shared across train, validation, and test. Positive BGC labels were assigned to genes matched to known clusters by antiSMASH KnownClusterBlast against MIBiG, while unmatched genes in the benchmark genomes were treated as negatives(Terlouw et al., 2023; Blin et al., 2023). Essential-gene, operon, and STRINGDB PPI benchmarks follow BacBench (Wiatrak et al., 2025b; Szklarczyk et al., 2023). The separate Fitness Browser analysis uses the February 2024 release of pooled randomly barcoded transposon sequencing (RB-TnSeq) experiments, which measure the fitness effects of gene disruption (Price & Deutschbauer, 2024). This analysis includes 179,946 genes with fitness records across 48 organisms. Table 3 summarizes the downstream datasets after filtering and split construction.

**Table 3:** Downstream datasets and probe settings. Split sizes give train/validation/test genome counts; zero-shot tasks report evaluation genomes in the test position and have no probe.

| Dataset | Metric | Split sizes | Loss |
| --- | --- | --- | --- |
| *Genome-level prediction* |  |  |  |
| Optimal growth temperature | Pearson | 3,713 / 795 / 795 | MSE |
| Doubling time | Pearson | 1,249 / 267 / 266 | MSE |
| Antibiotic resistance | Macro AUROC | 17,997 / 4,291 / 3,519 | BCE |
| MIC | Macro Pearson | 17,997 / 4,291 / 3,519 | MSE |
| Binary phenotypes | Macro AUROC | 17,080 / 3,660 / 3,660 | BCE |
| Mixed phenotypes | Macro F1 | 17,080 / 3,660 / 3,660 | MSE + CE |
| *Gene-level prediction* |  |  |  |
| BGC detection / category | AUPRC /<br>macro F1 | 555 / 61 / 153 | Weighted BCE<br>+ CE |
| Essential genes | AUPRC | 30 / 11 / 10 | BCE |
| Fitness Browser 2024 | AUROC | — / — / 48 | — |
| Operons | AUPRC | — / — / 11 | — |
| PPI | Spearman $\rho$ | — / — / 53 | — |

### A.2 Evaluation Protocols

All supervised evaluations use precomputed embeddings with frozen representation encoders. We use an MLP probe consisting of LayerNorm–Linear–GELU–Dropout–Linear, with dropout 0.1. Within each task, all representations use the same probe architecture and training hyperparameters, with input dimensions adjusted to the embedding size. Probes are trained with AdamW at a learning rate of 10^*−*3^ and weight decay of 0.01 for up to 100 epochs, with early stopping after 15 epochs without improvement in the task-specific validation criterion. For multi-output genome tasks, we average the applicable MSE, binary cross-entropy, or categorical cross-entropy losses over observed targets. Table 3 summarizes task-specific settings.

Zero-shot operon evaluation ranks gene pairs by embedding cosine similarity and reports AUPRC. For PPI, we treat each protein pair as unordered and count it once, compute the Spearman correlation between embedding cosine similarity and STRINGDB confidence scores within each genome, and report the unweighted mean across genomes. Appendix A.3 provides the protocol for zero-shot gene fitness evaluation.

### A.3 Gene Fitness Evaluation Details

We use the February 2024 Fitness Browser release, covering 46 bacteria and two archaea (Price & Deutschbauer, 2024). *For gene i*, we average its recorded fitness values as 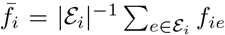, where ℰ_*i*_ is the set of recorded experiments, and assign the binary label 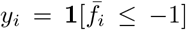. All remaining evaluated genes are negatives. Experiments receive equal weight, including repeated conditions, and missing values are not imputed. The target therefore summarizes fitness across the experimental panel available for each organism. Within the processed fitness-labeled subset, all genes in an organism have the same number of recorded experiments, ranging from 6 to 757 across organisms.

To compute representation sensitivity, VenusGenome replaces the target gene’s input protein embedding with the organism-wide mean input embedding, retaining its position, contig, and strand metadata. This mean is computed within the organism, not across the pretraining corpus. Let *N*_*i*_ contain up to five neighboring genes on either side of gene *i* on the same contig, excluding *i* itself. With ***h***_*j*_ and 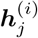 denoting the original and perturbed contextual states, respectively, the sensitivity score is

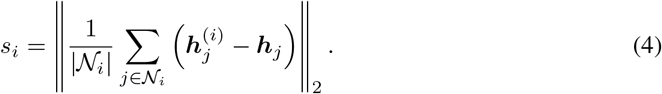

Neighborhoods are truncated at contig boundaries, and genes without eligible neighbors are excluded. We use the canonical 1B checkpoint with 4,096-gene windows and a 512-gene overlap.

The exploratory Evo2 baseline scores each standalone forward-strand CDS by its mean autoregressive log-likelihood. Higher representation sensitivity for VenusGenome and higher mean CDS log-likelihood for Evo2 indicate the positive class. We match model scores to experimental fitness measurements for each gene using its organism and gene identifiers. Both models are evaluated on the same 179,946 genes with finite scores and labels. We compute AUROC separately within each organism and report the median across organisms. All 48 organisms contain both classes, with 40–319 positive genes per organism.

### A.2 Model, Pretraining, and Extraction Details

Table 4 provides model cards for the three scales used in the scaling analysis; the main results use the 1B model. All models were trained on eight A800 GPUs with an effective batch of 256 genomes. Metagenome-assembled genomes account for 60% of training samples, split equally between MG-nify and SPIRE (30% each); the high-quality isolate collection accounts for the remaining 40%. We held out 2,000 genomes, evaluated every 5k steps with a fixed 15% mask (seed 2026), and saved checkpoints every 2.5k steps. Training and sampling seeds were 42. For downstream extraction, genomes longer than 4,096 genes are split into overlapping windows with 512-gene overlap, repeated gene states are averaged, and window-level global readouts are averaged. For genome-level evaluation, Evo2 and NT-v2 process genomic DNA in non-overlapping windows within each contig. Evo2 uses 8,192-bp windows and averages the last-token states from blocks.20, extracted in the forward direction. NT-v2 uses 6,000-bp windows, averages non-special token states within each window, and aggregates window representations weighted by their valid token counts. Neither pipeline subsamples windows; NT-v2 excludes fragments shorter than 50 bp.

**Table 4:** VenusGenome architecture and shared pretraining configuration.

| Configuration | 85M | 512M | 1B |
| --- | --- | --- | --- |
| **Architecture** |  |  |  |
| Protein representation | ESM2-35M, 480 dimensions |  |  |
| Encoder hidden size | 768 | 1,280 | 1,408 |
| Encoder layers | 12 | 26 | 44 |
| Encoder attention heads | 12 | 20 | 22 |
| Encoder FFN size | 3,072 | 5,120 | 5,632 |
| Decoder hidden size | 512 | 640 | 640 |
| Decoder layers | 8 | 8 | 8 |
| Decoder attention heads | 16 | 20 | 20 |
| Decoder FFN size | 2,048 | 2,560 | 2,560 |
| Transformer block | Pre-RMSNorm; GELU; dropout 0.1 |  |  |
| **Genome input and objective** |  |  |  |
| Genome context | 4,096 genes; 64 global tokens |  |  |
| Genomic structure | RoPE (random offset < 2,048); contig restriction; strand bias |  |  |
| Masking | 75% of genes |  |  |
| Reconstruction target | Global-mean-centered ESM2 embedding |  |  |
| Objective | 0.3 MSE + 0.7 cosine |  |  |
| **Optimization** |  |  |  |
| Optimizer | AdamW; $\beta = (0.9, 0.95)$; $\epsilon = 10^{-8}$; weight decay 0.01 | | |
| Learning-rate schedule | $10^{-4}$ peak; 5k-step warmup; cosine decay to $10^{-5}$ | | |
| Effective batch | 256 genomes on 8 GPUs |  |  |
| Training | 40k steps; BF16; gradient clipping 1.0 |  |  |

## B Additional Results

### B.1 Genome-Level Readout Comparison

We compare mean pooling of contextualized gene states with global-token aggregation using the same 1B mask-75% checkpoint at 40k steps. Both readouts are obtained from the same extraction run and evaluated with identical species-disjoint splits, labels, and probe configurations, using five paired probe seeds (1–5). Table 5 reports the mean and sample standard deviation using the metrics in Table 1.

**Table 5:** Mean pooling versus global-token aggregation on species-disjoint benchmarks.

| Readout | Temp. | Doubling | MIC | AMR | Binary phen. | Mixed phen. |
| --- | --- | --- | --- | --- | --- | --- |
| Mean pooling | 0.862<br>$\pm 0.001$ | **0.336**<br>$\pm 0.011$ | 0.270<br>$\pm 0.016$ | **0.682**<br>$\pm 0.034$ | 0.794<br>$\pm 0.004$ | **0.457**<br>$\pm 0.027$ |
| Global tokens | **0.877**<br>$\pm 0.002$ | 0.246<br>$\pm 0.015$ | **0.281**<br>$\pm 0.010$ | 0.672<br>$\pm 0.019$ | **0.797**<br>$\pm 0.003$ | 0.442<br>$\pm 0.023$ |

Table 5 shows that neither readout consistently outperforms the other. Global-token aggregation performs better on temperature prediction, whereas mean pooling has a larger advantage on doubling time. This suggests that the two readouts retain task-relevant information differently, with no single aggregation strategy preferred across all evaluated traits.

### B.2 Pretraining Loss Diagnostics

Figure 7 compares reconstruction loss under encoder scaling and matched architectural ablations. Larger encoders reach lower validation loss, consistent with the PPI scaling trend in Figure 3(a). For the 512M model, removing strand bias or global tokens increases validation loss, while removing contig-aware attention causes the largest degradation, mirroring the downstream ordering in Figure 3(c). At 40k steps, validation loss is 0.167 for the baseline, 0.277 without strand bias, 0.317 without global tokens, and 0.497 without contig-aware attention.

**Figure 7:**
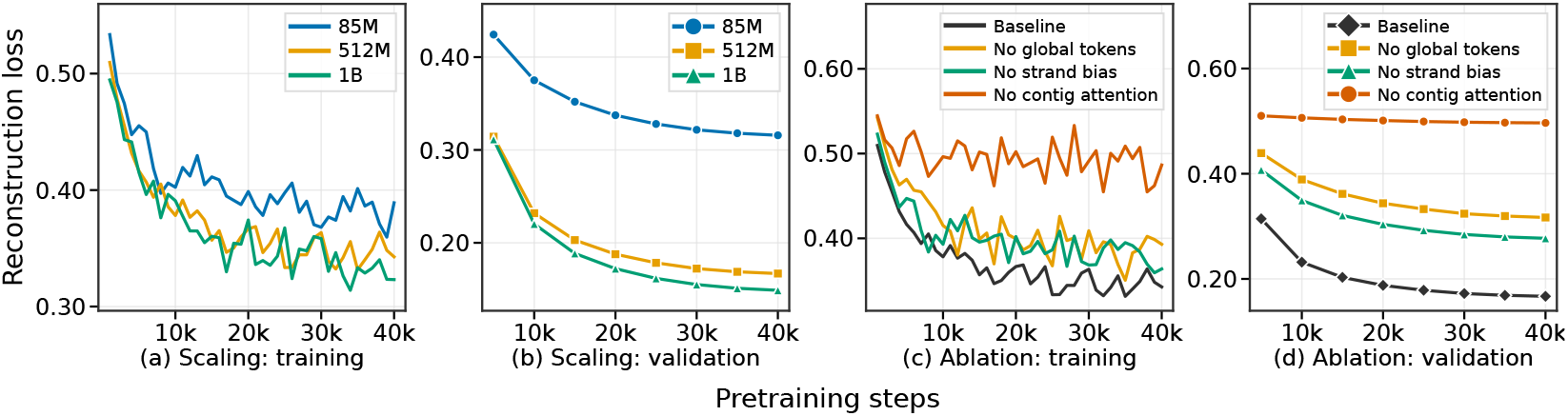
Pretraining reconstruction loss under encoder scaling and architectural ablations. (a,c) Training loss in 1000-step bins. (b,d) Validation loss evaluated every 5k steps with a shared fixed validation mask using 15% masked genes and seed 2026. Panels (a,b) compare 85M, 512M, and 1B mask-75% encoders; panels (c,d) compare the 512M mask-75% baseline with variants that remove global tokens, strand bias, or contig-aware attention.

### B.3 Mask Ratio and Pretraining Efficiency

We benchmark the computational cost of pretraining at different gene mask ratios using the 512M model. Higher masking ratios shorten the encoder input, while the decoder still processes all gene positions. On a single A800 GPU with batch size one and BF16 precision, we measure forward and backward computation through both the encoder and decoder, including the reconstruction loss, but excluding optimizer updates and distributed communication. The model, input, and contig-aware attention settings are held fixed. Reported reductions are averaged over four independent GPU runs, using the median of 20 timed steps after warm-up for runtime and peak allocated memory for memory usage.

Table 6 shows that the measured savings increase with sequence length. At 4,096 genes, increasing the mask ratio from 15% to 75% reduces forward–backward time by 65.2% and peak allocated memory by 51.3%. At 512 genes, runtime decreases by only 2.5% despite the shorter encoder input. Higher masking ratios thus reduce pretraining runtime and memory usage, with larger savings for longer genomic inputs in this benchmark.

**Table 6:** Measured pretraining efficiency of 75% masking (M75) relative to 15% masking (M15) for the 512M model. *L* denotes the number of valid genes; F+B denotes forward and backward passes. Encoder token counts include 64 global tokens.

| $L$ | Encoder tokens | | M75 reduction vs. M15 | |
| --- | --- | --- | --- | --- |
|  | M15 | M75 | F+B time | Peak memory |
| 512 | 499 | 192 | 2.5% | 4.1% |
| 1024 | 934 | 320 | 21.2% | 27.3% |
| 2048 | 1805 | 576 | 55.9% | 41.3% |
| 4096 | 3546 | 1088 | 65.2% | 51.3% |

## Notes

### Competing Interest Statement

The authors have declared no competing interest.

